# Kinship and social organization in Copper Age Europe. A cross-disciplinary analysis of archaeology, DNA, isotopes, and anthropology from two Bell Beaker cemeteries

**DOI:** 10.1101/863944

**Authors:** Karl-Göran Sjögren, Inigo Olalde, Sophie Carver, Morten E. Allentoft, Tim Knowles, Guus Kroonen, Alistair W.G. Pike, Peter Schröter, Keri A. Brown, Kate Robson-Brown, Richard J. Harrison, Francois Bertemes, David Reich, Kristian Kristiansen, Volker Heyd

**Author notes:** Corresponding authors: Volker Heyd, Department of Cultures – Archaeology, 00014 University of Helsinki, Finland,; Kristian Kristiansen, Department of Historical Studies, University of Gothenburg, 405 30 Gothenburg, Sweden.

## Abstract

We present a high-resolution cross-disciplinary analysis of kinship structure and social institutions in two Late Copper Age Bell Beaker culture cemeteries of South Germany containing 24 and 18 burials, of which 34 provided genetic information. By combining archaeological, anthropological, genetic and isotopic evidence we are able to document the internal kinship and residency structure of the cemeteries and the socially organizing principles of these local communities. The buried individuals represent four to six generations of two family groups, one nuclear family at the Alburg cemetery, and one seemingly more extended at Irlbach. While likely monogamous, they practiced exogamy, as six out of eight non-locals are women. Maternal genetic diversity is high with 23 different mitochondrial haplotypes from 34 individuals, whereas all males belong to one single Y-chromosome haplogroup without any detectable contribution from Y-chromosomes typical of the farmers who had been the sole inhabitants of the region hundreds of years before. This provides evidence for the society being patrilocal, perhaps as a way of protecting property among the male line, while in-marriage from many different places secured social and political networks and prevented inbreeding. We also find evidence that the communities practiced selection for which of their children (aged 0-14 years) received a proper burial, as buried juveniles were in all but one case boys, suggesting the priority of young males in the cemeteries. This is plausibly linked to the exchange of foster children as part of an expansionist kinship system which is well attested from later Indo-European-speaking cultural groups.

## Introduction

Recent genetic research has made it clear that the third millennium BC was a period of large-scale migrations from the Caspian-Pontic steppe towards central and, later, western Europe, leading first to the formation of the Corded Ware and then the Bell Beaker complexes [1, 2, 3]. This is also evidenced in a shared burial ritual in Central Europe, characterized by individual burials and strict differentiation between males and females in the orientation of the body [4]. The genetic admixture that resulted from these migrations still characterizes modern European populations, just as it is very likely that predecessors of one or several Indo-European languages spoken in Europe today were carried by these migrations [5, 6].

However, it is still not well understood how it was possible for these populations to establish and maintain their cultural, social and linguistic coherence over time. For the Corded Ware complex it has been suggested that initial migrations were dominated by males, who married in women probably from residing Neolithic populations [6, 7, 8, 9], although at present it is debated whether the genetic evidence for male-dominated migrations contributing to these groups is compelling [10, 11]. There is also evidence that the individual groups most likely practiced patrilocality and exogamy at a community level [12, 7].

We wanted to test if such a pattern of kinship, patrilocal residence and exogamous marriage persisted for the following Bell Beaker culture in Central Europe, which has been partly suggested by recent research on Bell Beaker and Early Bronze Age burial communities from the Lech valley around the city of Augsburg in Bavaria [13; 14]. We also wanted to test a proposition from Knipper et al. [13] that such marriage pattern would lead to increasing gene pool diversification. Here, we propose that such a degree varied with the complexity of social alliances. Finally, we wish to test if male lineages were maintained through time. In the concluding section, we discuss which social mechanisms and institutions would support such a long-term genetic and possibly linguistic stability.

Our reference point to assess these hypotheses are findings coming from two cemeteries of the late Bell Beaker culture of South Germany [15], both located close to the Danube river, only 17 kilometers apart from each other, and roughly belonging to the same chronological horizon [Fig 1). Both cemeteries, Irlbach (IRL; county of Straubing-Bogen) and Alburg (ALB; -Lerchenhaid; city of Straubing) were entirely excavated during rescue excavations in the 1980s and are documented as 24 and 18 graves, respectively.

**Fig 1.**
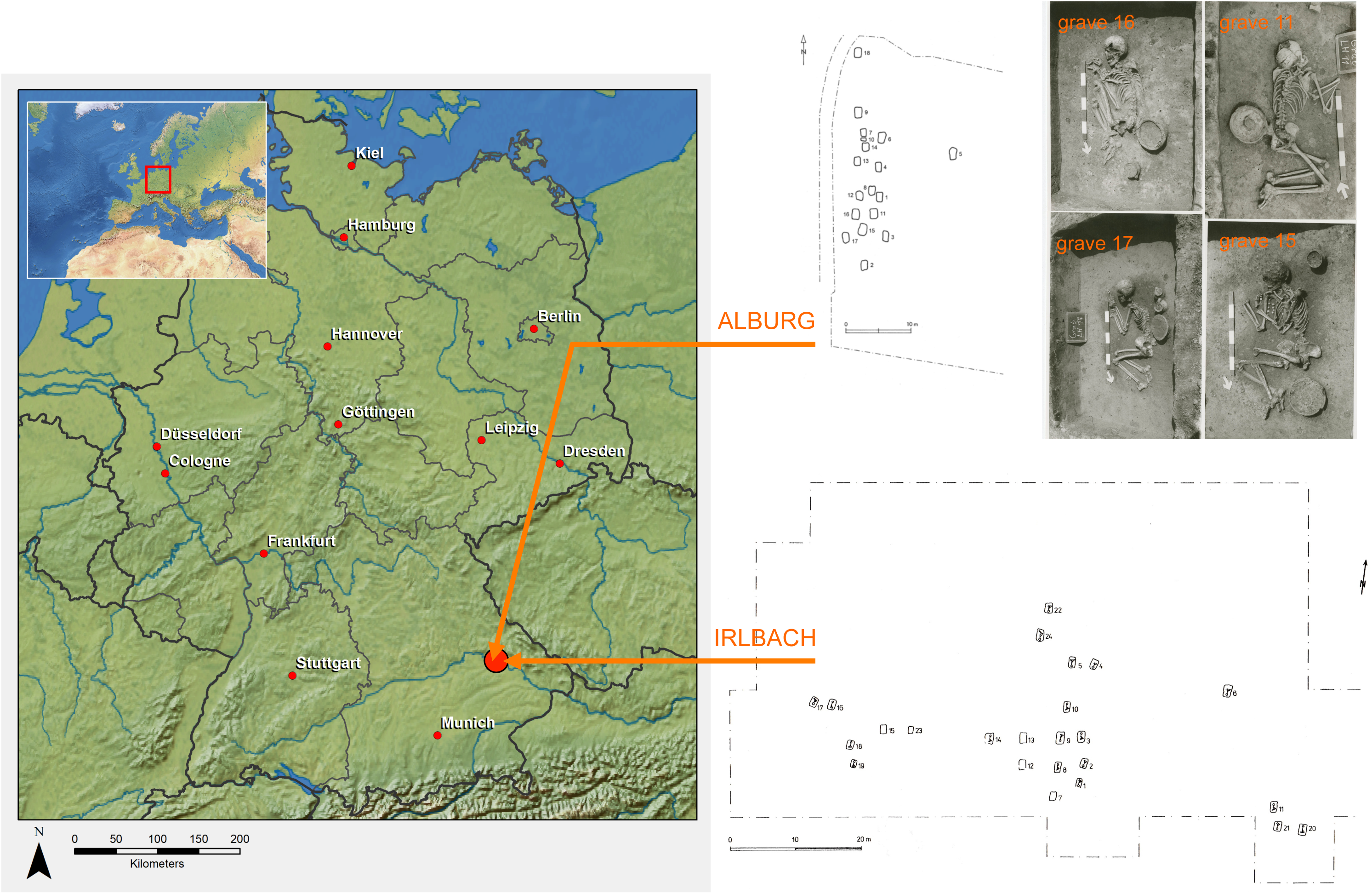
Location and plans of the two late Bell Beaker culture cemeteries of Irlbach and Alburg (Straubing, Bavaria, Germany); the graves nos. 11, 15, 16 and 17 from the Alburg cemetery are shown as examples.

Both cemeteries have received a detailed archaeological assessment, highlighting funerary customs, material culture and chronological sequences. They have both been fully analyzed bio-anthropologically, subjected to multi-isotopic measurements of tooth enamel, and were part of the recent Europe-wide Beaker Phenomenon ancient DNA project [3] in which context their sex, mitochondrial and Y-chromosome haplogroups, ancestry, and genetic kinship were established. Here we take the genomic analyses one step further to investigate kinship between the individuals in detail.

## Materials and Methods

### Archaeological Background

The cemeteries of Alburg (Lerchenhaid-Spedition Häring, City of Straubing, Lower Bavaria, Germany) and Irlbach (County of Straubing-Bogen, Lower Bavaria, Germany) were excavated in rescue excavations, the one in 1982 by the State Heritage Office in Landshut and the other in 1987-89 by the county archaeologist of Straubing-Bogen, Karl Böhm [15, vol. 2: 64-65; 72-73; 3, SI_2: 37-41]. Grave documentations are deposited in the County Archaeology Office in Bogen and in the Bavarian State Office for Heritage in Regensburg, respectively. All finds from both cemeteries are in the Gäubodenmuseum in Straubing. Human remains are stored in the premises of the State Collection for Anthropology and Palaeoanatomy in Munich. Both cemeteries, only being 17 kilometres apart from each other, have a similar location in that graves are dug into the löss soil cover of the lower terrace of the right Danube riverbank. Contemporary settlement sites, potentially belonging to these cemeteries, are unknown. Cultural attribution is based on archaeological criteria, such as grave goods and burial customs.

### Bio-anthropological examinations of skeletons

Anthropological sex was established according to Brothwell’s [16] pelvic measurements, as well as the scales (from one to five) of cranial sexing from White and Folkens [17]. In addition, where the above two methods were not possible, sex has been ascertained through metrics as seen in Brothwell [16], Bass [18], and Chamberlain [19]. Age was established through epiphyseal fusion in Scheuer and Black [20], cranial suture closure (where possible) from Meindl and Lovejoy [21], dental development phases from White and Folkens [17], and dental attrition in Brothwell [16]. Non-metric traits were scored as present, absent, or not observable. Cranial and post-cranial non-metric traits were taken from Berry and Berry [22], Brothwell [16], Buikstra and Ubelaker [23], Mann, Hunt, and Lozanoff [24] and Hauser and De Stefano [25]. Metrics have been measured, or scored as absent, and stature estimations have been made, where possible, according to Trotter [26]. Preservation has been assessed on a scale of good, medium, or poor, and an approximation of percentage of remains present has been recorded. Surface erosion of the bones was assessed according to McKinley [27].

### Strontium and Oxygen Isotope Analyses

^87^Sr/^86^Sr isotope analysis of the Irlbach cemetery enamel samples were taken from previous publications [28, 29]; see for an overview also [30]. A detailed list of the sampled teeth is provided on Fig 7 of [31]. Samples for Alburg cemetery’s ^87^Sr/^86^Sr and both cemeteries’ δ^18^O isotope analysis were taken as follows. The enamel surface of an intact tooth was first cleaned using a dental burr and hand drill. Two wedges of enamel and dentine (ca. 0.5mm wide, 1mm deep) representing the complete growth axis of the enamel were removed using a flexible diamond impregnated dental disc. Any dentine adhering to the enamel sections was then removed using a dental burr, and the remaining enamel sample cleaned in an ultrasonic bath. For Sr isotope analysis, the whole enamel section was dissolved in 3ml 7N HNO3. Any detritus was removed by centrifuging, and the supernatant was dried and redissolved in 3N HNO3. An aliquot of this solution was removed, representing 3mg of solid enamel (containing approximately 100-300 ng of Sr), and made up to 0.5ml 3N HNO3 to be loaded onto ion exchange columns. The strontium was separated using standard ion exchange chromatography using 70μl of Eichrom Sr spec resin (50-100 μm). Samples were loaded in to 0.5ml 3N HNO3 and washed with 4ml 3N HNO3. Strontium was eluted in 1.5ml MilliQ water. The elutant was dried down and loaded using a few μl 10% HNO3 onto rhenium filaments preconditioned with 1 μl TaCl5 solution and 1 μl 10% H3PO4. Isotope ratios were measured on a ThermoFinnegan Triton Thermal Ionization Mass Spectrometer in the Department of Earth Sciences of the University of Bristol, UK. The data is corrected for mass fractionation using a ^86^Sr/^88^Sr value of 0.1194 and an exponential fractionation law. ^87^Rb is subtracted using the measurement of ^85^Rb and a ^85^Rb/^87^Rb value of 2.59265 [32]. Data is corrected to NBS 987 using a value of 0.710248. The typical precision for ^87^Sr/^88^Sr achieved for a tooth sample using this method is ±0.00001.

For oxygen isotope analysis of both cemeteries, the cleaned enamel sample was ground to a power in a hand mortar. The oxygen isotopes were taken from the structural carbonate fraction of tooth enamel (δ^18^Oc) and were measured relative to Pee Dee Belemnite (PDB) by Peter Ditchfield, Research Laboratory for Archaeology and the History of Art, University of Oxford, UK, on an Isoprime Dual-Inlet mass-spectrometer connected to a Gilson auto-sampler using Oxford standard carbonate procedures.

### Ancient DNA Analyses

We extracted DNA from four newly reported individuals from Alburg and Irlbach and generated double-stranded DNA libraries following the same protocols as in [3]. We also generated additional DNA libraries from 14 individuals included in [3]. All the libraries were subjected to a partial uracil-DNA-glycosylase (UDG-half) treatment to reduce the effects of post-mortem cytosine deamination [33]. Libraries were captured with probes overlapping 1,233,013 SNPs (‘1240k capture’) and the mitochondrial genome [34, 2], and sequenced on Illumina NextSeq500 instrument with 2×76 cycles and 2×7 cycles to allow the indices to be read. Sequencing reads were processed bioinformatically as in [3]. For the four newly reported samples mitochondrial haplotypes were called using Haplogrep2 [35]. For the 14 individuals with additional libraries, new sequencing data were merged with data from [3], and Y-chromosome calls from [3] were updated accordingly (Table 1).

**Table 1.** Overview table of Irlbach and Alburg grave number, osteological and genetic sex, age group at death, burial position, archaeological dating/period, grave goods, ^87^Sr/^86^Sr enamel and bone values, δ^18^Oc values, mtDNA haptotype and Y DNA haplogroup.

| Site | Grave | Osteo-logical sex | Genetic sex | Age group | Position | Arch period | 87Sr/86Sr enamel | 87Sr/86Sr bone | mtDNA haplo | YDNA haplo |
| --- | --- | --- | --- | --- | --- | --- | --- | --- | --- | --- |
| Irlbach | 1 | F | F | Adult | Right side | B1 | 0.71028 | 0.71092 | U5b2c |  |
| Irlbach | 2 | F | F | Young adult | Right side | B1 |  |  | X2c1 |  |
| Irlbach | 3 | M | M | Adult | Left side | B1 | 0.70963 | 0.70952 | T2b+152 | R1b1a1a |
| Irlbach | 4 |  | F | Inf I | Right side | B1 | 0.70971 |  | H5a1 |  |
| Irlbach | 5 |  | F | Inf II | Right side | A2 | 0.7095 |  | U5a1a2b |  |
| Irlbach | 6 | F | F | Mature adult | Right side | B2 | 0.71241 | 0.70913 | H5a1 |  |
| Irlbach | 7 |  | undet | Inf I |  | B1 | 0.70964 |  | T2g2 |  |
| Irlbach | 8 |  | M | Inf II | Left side | B1 | 0.71006 |  | T2b+152 | R1b1a1a2<br>(M269) |
| Irlbach | 9 | F | F | Mature adult | Right side | B1 | 0.70955 | 0.70932 | T2b+152 |  |
| Irlbach | 10 | M | M | Adult | Left side | A2 | 0.70938 | 0.70938 | K1a4b | R1b1a1a2a1a2b1<br>(M269) |
| Irlbach | 11 |  | M | Inf II | Left side | B1 | 0.71096 |  | J1c+489+1598+3504<br>+12477+16188+16189 | R1b1a1a2a1a2b1 |
| Irlbach | 12 |  |  | Young adult | Left side? | B2 |  |  |  |  |
| Irlbach | 13 |  |  | Mature adult | Right side |  |  |  |  |  |
| Irlbach | 14 | M | M | Adult | Left side | B1 | 0.7094 | 0.70914 | T1a1 | R1b1a1a2 |
| Irlbach | 15 |  |  | Juvenile |  | B1 |  |  |  |  |
| Irlbach | 16 | M | M | Adult | Left side | B2 | 0.71158 | 0.70933 | K1b1b1 | R1b1a1a2 |
| Irlbach | 17 | F | F | Mature adult | Left side<br>"Rückenhocker" | B1 |  |  | W5 |  |
| Irlbach | 18 | M |  | Mature adult | Left side | B1 |  |  |  |  |
| Irlbach | 19 |  | M | Inf I | Left side |  |  |  | U5a2a+16294 | R1b |
| Irlbach | 20 | M | M | Adult | Left side | B2 | 0.70981 | 0.70922 | J1c | R1b1a1a2a1a2b1 |
| Irlbach | 21 | F | undet | Juvenile | Right side | B2 | 0.71013 |  | HV6 |  |
| Irlbach | 22 | F | F | Adult | Right side | B1 | 0.70972 | 0.70929 | T1a1 |  |
| Irlbach | 23 |  |  |  |  |  |  |  |  |  |
| Irlbach | 24 |  |  |  | Right side |  |  |  |  |  |
| Alburg | 1 |  | M | juvenile | Left side |  | 0.70936 |  | H1e1a | R1b1a |
| Alburg | 2 |  | M | Adult | Left side |  | 0.70951 |  | H1e1a | R1b1a1a2a1a |
| Alburg | 3 |  | M | Adult | Left side |  | 0.70986 |  | U4d1 | R1b1a1a2a1a2b1 |
| Alburg | 4 |  | F | Adult | Right side | B1 | 0.70986 |  | H1+10410+16193+16286 |  |
| Alburg | 5 |  |  |  |  |  |  |  |  |  |
| Alburg | 6 |  | F | Adult | Right side | B2 | 0.70993 |  | H1+10410+16193+16286 |  |
| Alburg | 7 |  | undet | Inf II | Left side | B1 | 0.70981 |  | H1+10410+16193+16286 |  |
| Alburg | 8 |  | undet | Adult | Right side | B1 | 0.7107 |  | H+16129 |  |
| Alburg | 9 |  | F | Adult | Right side | A2 | 0.71067 |  | T2f |  |
| Alburg | 10a |  | undet | Inf I |  | B1 |  |  | V |  |
| Alburg | 10b |  |  | Inf I |  | B1 |  |  |  |  |
| Alburg | 11 |  | M | Juvenile | Left side | B1 | 0.70981 |  | H+16129 | R1b1a1a2 |
| Alburg | 12 |  | M | Inf II | Left side | B1 | 0.70952 |  | T2f | R1b1a1a2a1a2b1 |
| Alburg | 13 |  | M | Adult | Left side | A2 |  |  | H1e1a | R1b1a1a2a1a2 |
| Alburg | 14 |  | F | juvenile/Adult | Left side | B1 | 0.70956 |  | U5b3 |  |
| Alburg | 15 |  | F | Adult | Right side | A2 | 0.71014 |  | H10e |  |
| Alburg | 16 |  | F | Adult | Right side | A2 | 0.7174 |  | I3a |  |
| Alburg | 17 |  | undet | Adult | Right side | B2 | 0.70925 |  | H1+10410+16193+16286 |  |
| Alburg | 18 |  |  |  | Cremation |  |  |  |  |  |

We determined genetic sex based on the ratio of Y chromosome to sum of X and Y chromosome sequences [36] and relatedness coefficients using the method in [37] with nucleotide mismatch rate of 0.127. For Principal Component Analysis (PCA), a random allele was sampled for each ancient individual at each of the 591,642 SNP positions included in the analysis, removing the first and last two nucleotides of the sequences to avoid the effects of DNA damage. Principal components were computed on 989 present-day West Eurasians genotyped on the Human Origins Array, using the ‘smartpca’ program in EIGENSOFT [38]. Individuals from Alburg and Irlbach, as well as other previously published ancient individuals [39, 1, 40, 41, 42, 43, 44, 3] from relevant populations, were projected onto the components computed on the present-day individuals with lsqproject:YES and shrinkmode:YES.

## Results

### Archaeology and Bio-Anthropology

The 24 graves make Irlbach the largest Bell Beaker culture cemetery in South Germany discovered to date. However most graves have been damaged by ploughing, and likely several more were completely destroyed prior to the excavations. The cemetery might originally have included ∼30 graves, arranged in three west to east groups, on an overall area of ∼60 (W-E) × 30 (N-S) m. Of these groups, the western part yields six, the central part 14, and the eastern part three graves plus one more isolated grave (IRL_6). The Alburg graveyard, in contrast, is perfectly preserved and appears denser in its occupation with 18 graves covering an area of ∼10 × 30 m. Almost all graves, with grave pits sized up to 1,4 × 0,8 m, are laying in long rows, oriented north-south. Only grave ALB_5 is off one of these rows, and it could not be established whether this really belongs to the cemetery.

Out of the 41 graves that it has been possible to document, all are individual inhumation graves in often quite shallow burial pits, mainly orientated north-south, and often arranged in lines of graves or clustered together. Exceptions are grave ALB 18 which is a cremation; ALB 10 yielding two infant I (0-7 years) children (maybe neonate twins); and IRL 2 of a ‘young adult’ woman and an infant I child. Most burials are furnished with pottery, predominantly one or two cups and/or plate/bowl with six graves from Irlbach having single animal bones as remains of original food offerings in the latter. Only ALB 9 had a decorated beaker vessel, of the type that gives its name to the Bell Beaker complex. Beyond ceramics, male graves contained occasional flint arrowheads, deer teeth, and decorated tusk/bone pendants usually associated with hunting (four men in Irlbach and one, ALB 3, in Alburg). Female graves contained a series of small V-bored bone/antler buttons. While three burials in Irlbach contained only a few, the Alburg cemetery stands out due to six graves yielding many, amongst these 29 pieces in grave 6 alone and 22 pieces in grave 15, here laid out “in a U-formed line from the clavicle to the lower departure of the sternum and then upwards again to the other clavicle”, most of them with the perforated side facing upward. Both sexes show few signs of social differentiation based on the presence of prestige objects; there is only one small copper dagger but no wristguards, or artefacts made of gold, silver or amber, which could otherwise be linked with status.

The material culture forms the basis of a chronological sequencing of the burials in which characteristic groups of equipment and pottery lead to the definition of the relative phases A2b, B1 and B2, each likely comprising a few generations [15, 45]. Both cemeteries show a similar sequence, spanning from phase A2b to B2, with most of the graves in phase B1. In archaeological terms, they may therefore be regarded as mostly contemporary, with the graves IRL 5 and IRL 10 being the earliest in Irlbach. The four graves of the eastern grave group (IRL 6, IRL 11, IRL 20 and IRL 21) are the latest interred, also representing the latest Bell Beaker stage in Bavaria.

In Alburg, graves ALB 18, ALB 9, ALB 13, ALB 16 and ALB 2 are the earliest and arranged in an initial north-south orientated line of burials. Grave ALB 6 and ALB 17 are seemingly the latest. Both cemeteries were in use for a period of more than ∼100 years. To secure the chronology, four radiocarbon datings were performed. Despite long 2σ-calibration spans in the second half of the third millennium due to wiggles in the calibration curve, these generally support a date of ∼2300-2150 cal BC, consistent with other middle to late Bell Beaker cemeteries in Bavaria, but are too few and imprecise to improve the estimates of the use-life of the cemeteries beyond the relative dating based on material culture and genetics.

Southern German Bell Beaker culture people practiced a gender differentiated burial custom in which most females lay crouched on their right sides, with heads to the south, and most male individuals lay on their left with heads to the north (examples are shown in Fig 1). Both bio-anthropological and genetic sexing (Fig 2) confirms a broad adherence to this custom. In each cemetery we however find exceptions: Grave IRL 17 is that of a mature woman whose legs probably lay on the wrong side (the left instead of the right); moreover she might even have been positioned supine with flexed legs. ALB 14 turned out to be genetically female despite having a male body position.

**Fig 2.**
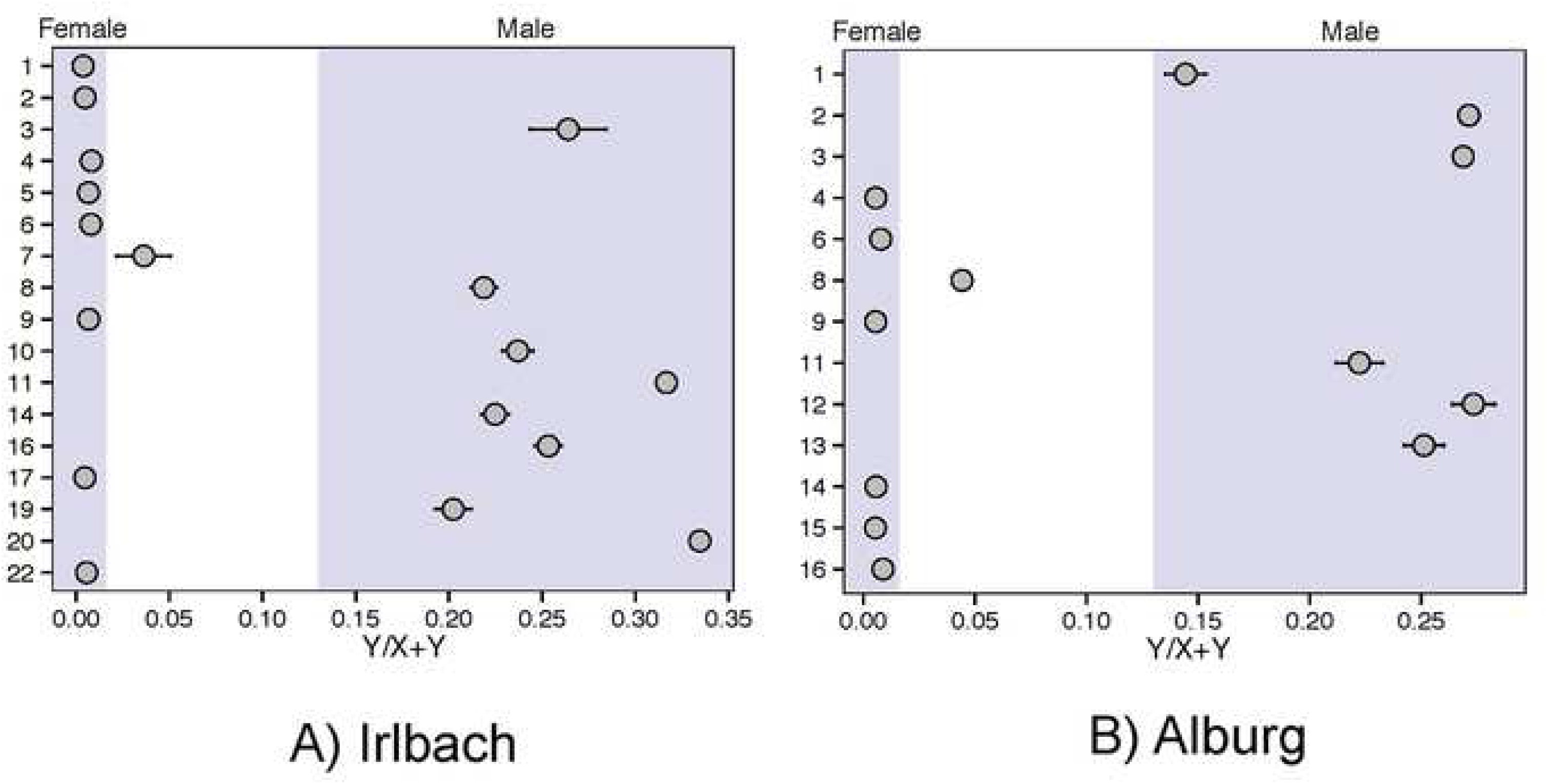
Genetic sexing results of the A) Irlbach and B) Alburg cemeteries.

Sexes are well balanced in both cemeteries: Irlbach includes 11 females and nine males, while Alburg contains seven females and eight males. In Irlbach, 14 individuals reached adulthood (six men and seven women, with one of unknown sex) and in Alburg ten (three men and seven women). Irlbach has seven infant I (0-7 years) and II (7-14 years) burials (at least two girls and three boys), and two juveniles (one girl, IRL 21; the other of unknown sex). Alburg has five infant I and II burials, two boys and at least one girl, but likewise only two juveniles (both boys). Children of the infant age group are thus underrepresented in what one would expect for pre-industrial societies with high child mortality [46, 47]. This suggests that the communities practiced a social system that selected children to be allowed a proper burial in the communal cemeteries. Such a system may also have been in place for adolescents as the Fig for both cemeteries combined seems unbalanced, favoring boys for burial (seven versus four girls).

The burials in both cemeteries have given no anthropological evidence for the causes of death. There are only few pathologies, minimal evidence for malnutrition, and only one case of interpersonal violence, represented by the man in IRL 14. He displays a remodeled right radius and ulna break just above their distal joint surfaces. Being well above average stature and one of the tallest men in the series, he is also the only individual in both cemeteries having a copper dagger, originally placed at the right radius/ulna. Another copper object may have originally been given to grave IRL 22, however it was removed already in antiquity. Grave IRL 20 was also intentionally disturbed.

The non-metric trait of a *septal aperture*, the incomplete fusion of the distal joint surface of the humerus, is displayed in the skeletons from IRL 3, IRL 14, IRL 21 and IRL 22, consisting of two men and two women, one of whom is a juvenile, 15-16 years old. This trait is represented with only 6% in today’s general population [48]. Compared to the altogether nine individuals with at least one fused distal joint surface of the humerus in the collection, it is disproportionally represented in the Irlbach skeletal series. It is probable that the number of these traits found in Irlbach is a result of hereditary inheritance and thus kinship.

### Ancient DNA

We possess genetic data for 18 graves from Irlbach and 16 graves from Alburg (Table 1). This set can be divided into 1) Y-chromosome haplogroups; 2) Mitochondrial DNA (mtDNA) haplotypes; and 3) Hundreds of thousands of autosomal markers allowing high-resolution ancestry inferences and kinship analysis. Four individuals (three from Alburg and one from Irlbach) with only mtDNA information are newly reported in this study, and we generated additional DNA libraries on 14 individuals (SI Table) reported in a recent study [3]. The new data are released on the Reich laboratory website as well as at the European Nucleotide Archive at accession number xxx.

All the male burials with sufficient data belong to a single Y-chromosome lineage, R1b-M269, which is the major lineage associated with the arrival of Steppe ancestry in western Europe after 2500 BC. In the preceding and partly contemporary Corded Ware populations of central Europe, another Y-haplogroup dominated, R1a, although R1b also occurs albeit in small numbers [1]. For individuals for whom we can determine the R1b-M269 subtype, we found that all had the derived allele for the R1b-S116/P312 polymorphism, which defines the dominant subtype in central and western Europe today [3]. This represents an extraordinary uniformity along the male line, practically linking all men in both cemeteries and in fact the vast majority of Central European Bell Beaker culture men who are also R1b-S116/P312 positive [3]. However, given that this lineage likely arose several centuries earlier, this uniformity does not necessarily imply a very close paternal relationship between the males of these two communities.

In stark contrast to the patterns of Y chromosome variation, the 18 individuals in the Irlbach graves have 14 different mitochondrial haplotypes and the 16 individuals in Alburg still share nine, showing an extreme diversity of maternal lines (Fig 3A-B). This suggests the possibility of widespread, probably institutionalised, exogamic marriage pattern incorporating for generations women from various backgrounds into burial communities and netting them together in extended kin-groups. Interestingly, none of the haplotypes is shared between our two cemeteries. This speaks for our two burial groups belonging to two different wider kin-groups despite their spatial proximity. Compared to the only other contemporary set recently made available [13], i.e. three Bell Beaker culture burial groups and two single graves around Augsburg, ca 200 kilometres away, with their 16 different haplotypes out of 19 positively tested burials, both Augsburg and Irlbach show a similar mitochrondrial diversity, while it is lower at Alburg (Fig 3C).

**Fig 3.**
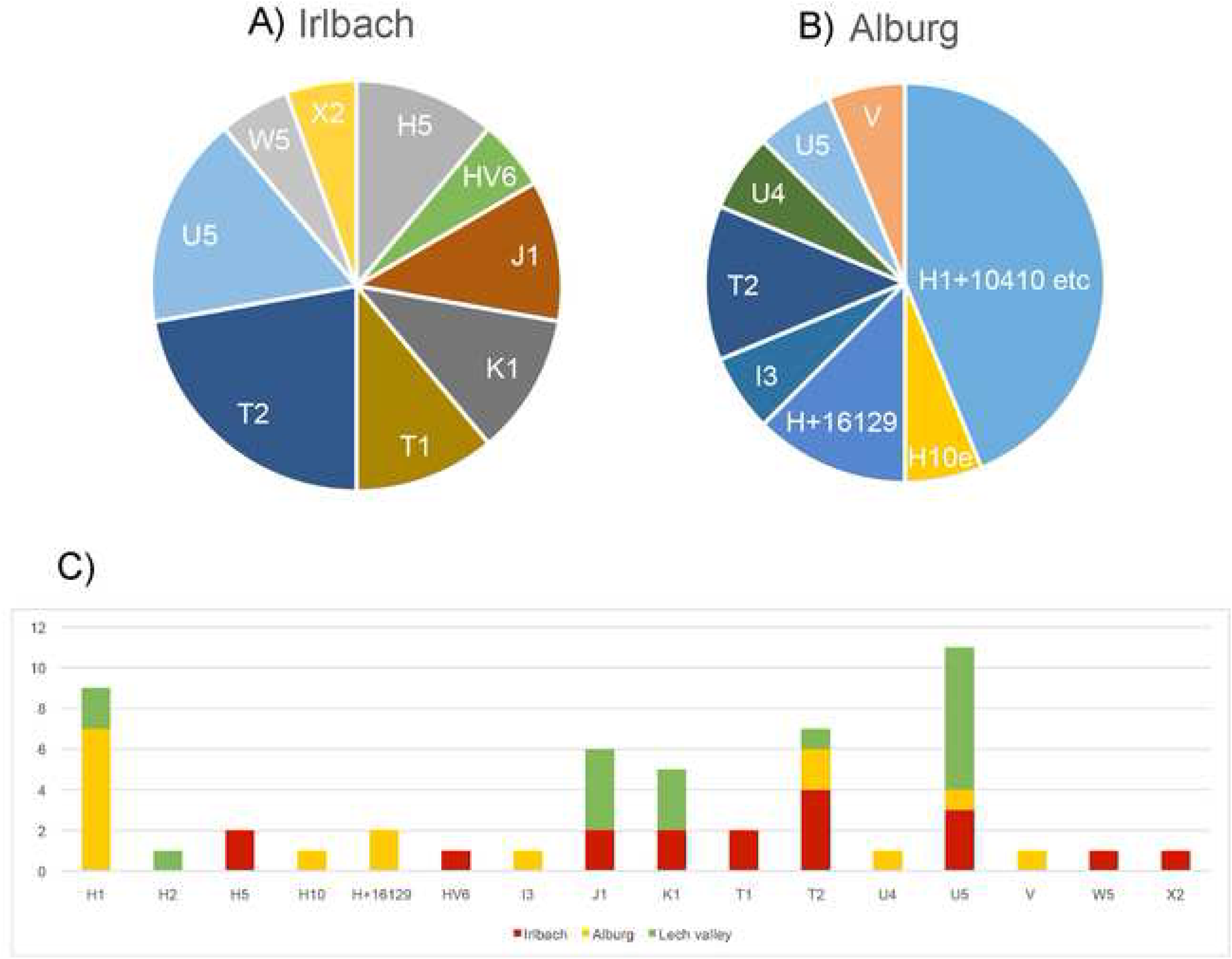
Pie chart of mtDNA haplotype distribution of A) Irlbach and B) Alburg, in comparison C) with the Bell Beaker cemeteries around Augsburg.

The genome-wide ancestry of 16 individuals from Irlbach and 13 from Alburg is illustrated in a principal component analysis (PCA) projecting the ancient samples onto the genetic variation in a set of west Eurasian present-day populations (grey dots), with previously published (pale yellow) ancient samples (Fig 4). The results show that individuals from both cemeteries ranged along the cline determined by Bronze Age Steppe and European Neolithic ancestries, with IRL 9, IRL 10 and IRL 16, and ALB 14 and ALB 16 having closer affinity to Steppe/Corded Ware populations, while IRL 4 and IRL 14, and ALB 4, ALB 6, ALB 9, and ALB 12 are more leaning towards much earlier established (= pre-Yamnaya) European Early and Middle Neolithic farmers. This picture is basically identical to the later situation around Augsburg [14].

**Fig 4.**
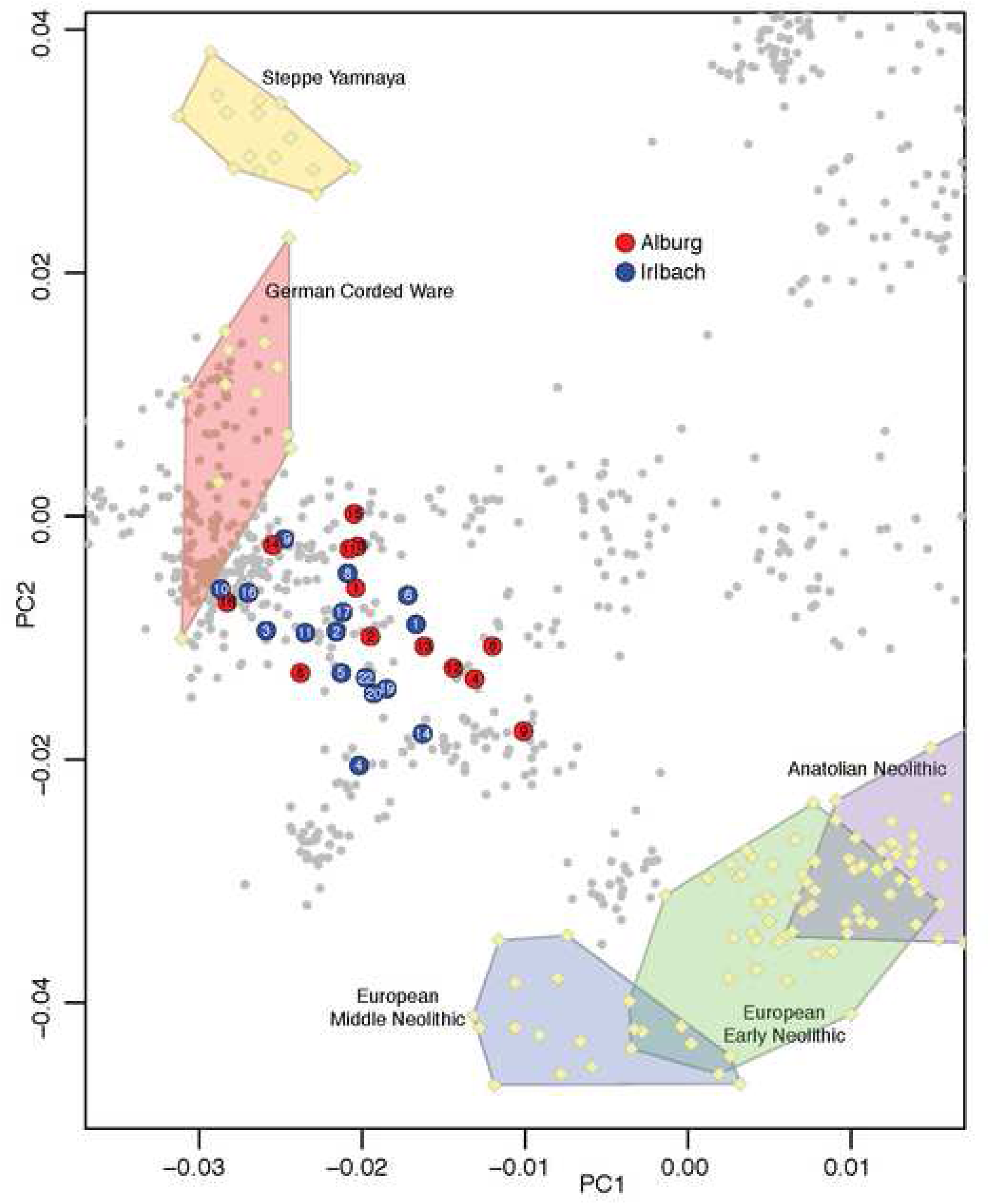
Principal Component Analysis using ∼600000 autosomal genetic markers on 990 present-day West Eurasians (shown as grey circles). Ancient individuals are projected onto the first two principal components computed on the present-day individuals, to avoid the effects of ancient DNA damage.

Using genome-wide data, we determined intra-group kinship (Fig 5), identifying 1st-degree-relationships among individuals from the IRL 3–8–9, IRL 11–20 and IRL 14–22, as well as ALB 4–6, ALB 9–12, ALB 2–13, and likely ALB 1–2 and ALB 1–13. In combination with sex/age information, grave location and position in the chronological sequence (older/younger), further conclusions can be drawn: IRL 3, IRL 8 and IRL 9 lay in the centre of the cemetery and are likely earlier than IRL 11, IRL 14, IRL 20 and IRL 22. IRL 8 and IRL 9 are either siblings or mother (died aged 50+) and juvenile son (died aged 10-11). The adult man (died aged 30-40) in IRL 3 is also a 1st-degree relative of the individuals in IRL 8 and IRL9. As they share the same mitochondrial haplotype, individuals in these three graves, placed next to each other, are likely those of a mother and her two male children. Less likely seems the combination of two brothers and their sister. IRL 20 is the father of IRL 11, who is a juvenile boy with a different mtDNA haplotype. IRL 14, an adult man aged 40-45 years at death, is either a sibling or the son of the adult woman IRL 22, passed away at age 24-25.

**Fig 5.**
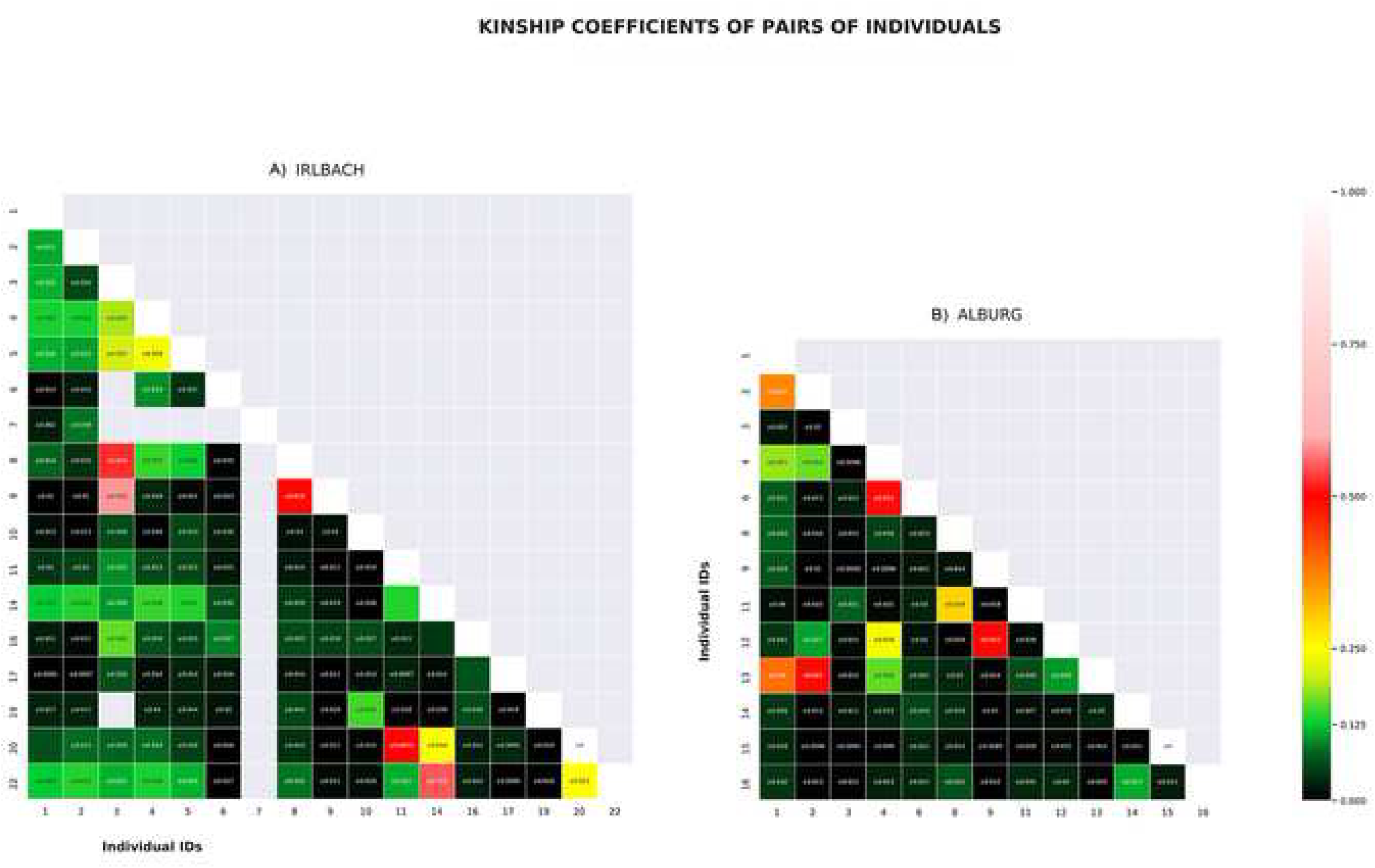
Genetic intra-group kinship results of the A) Irlbach and B) Alburg cemeteries.

ALB 4 is likely the mother of the woman in ALB 6. The adult woman in ALB 9 is the mother of the male child ALB 12. ALB 2 and ALB 13, both adult men at death, are brothers as they share the same haplotype of H1e1a. This haplotype is also shared by ALB 1, a juvenile boy buried without pottery like ALB 2, and located slightly offset in between 2 and 13. He also shares a high kinship coefficient with both other brothers. A scenario that best fits these relationships, and is consistent with the temporal succession of burials, would see the individuals in ALB 1–2–13 being first generation brothers.

We also detected 2nd and 3rd-degree-relationships and more distant kinship. In Irlbach these are 2nd-degree kin pairs in IRL 14–20 and IRL 20–22 and 3rd-degree kin pairs in IRL 11–14 and IRL 11–22. This link shows that all four graves are closely related, consistent with the phenotype of septal aperture of their humeri, with the most likely scenario being that the individual IRL 20 is not only the father of IRL 11 but also the nephew of both the adult woman in IRL 22 and the adult man in IRL 14. Despite sharing the same mitochondrial H5a1 haplotype with an exact match, IRL 4 and IRL 6 are not first or 2nd-degree relatives, but they could be 3rd-degree relatives. IRL 1–2–4–5 are likely 3rd-degree relatives, with the exception of IRL 4–5 who are more likely 2nd-degree relatives. IRL 1, IRL 2, IRL 4, and IRL 5 are also 3rd-degree or more distant relatives of IRL 14–22–11–20. Finally, brothers IRL 3–8 are likely 3rd-degree or more distant relatives of IRL 1–2–4–5–14–22–11–20.

In Alburg, ALB 4–12 are 2nd-degree relatives and ALB 8–11 sharing the same mtDNA H+16129 haplotype are likely also 2nd-degree kin. The likely brothers in the ALB 1–2–13 are equally related to ALB 4 as 2nd/3rd-degree kin, with one possibility being niece and paternal uncles relationships. ALB 1–2–13 are also 3rd-degree relatives of ALB 12. ALB 14–16 are likely 3rd-degree relatives. Finally, ALB 7 and ALB 17 (both with only mtDNA data) share the same mtDNA haplotype (H1+10410+16193+16286) with ALB 4–6, suggesting a close maternal relationship.

The adult woman in ALB 4 could then be the daughter of an unsampled brother of ALB 1–2–13. Given that grave ALB 4 is related with the male child in ALB 12 but not with his mother ALB 9, ALB 4 would be the paternal aunt/niece of ALB 12. While we cannot see her parents amongst the burials, this woman ALB 4 appears like the kinship ‘hinge’ for the first two and the last generation(s), her own daughter being in ALB 6 and her infant children or grandchildren perhaps in ALB 7 and ALB 17.

These genetic connections make it likely that we have close-knit kin-groups in both cemeteries, with the 10 adult individuals in Alburg (Fig 6) forming a single nuclear family over several generations, likely ∼four to five. In Irlbach (Fig 7), the six individuals of the western burial group seem unrelated to each other and to the central group within the limits of our sampling and resolution. One can therefore estimate the existence of one, more extended family group as the eastern three burials of Irlbach, and the isolated IRL 6, are genetically linked to the main group. The duration of use of the cemetery as a burial place might also encompass ∼five to six generations however this remains difficult to calculate due to the mentioned destruction due to ploughing.

**Fig 6.**
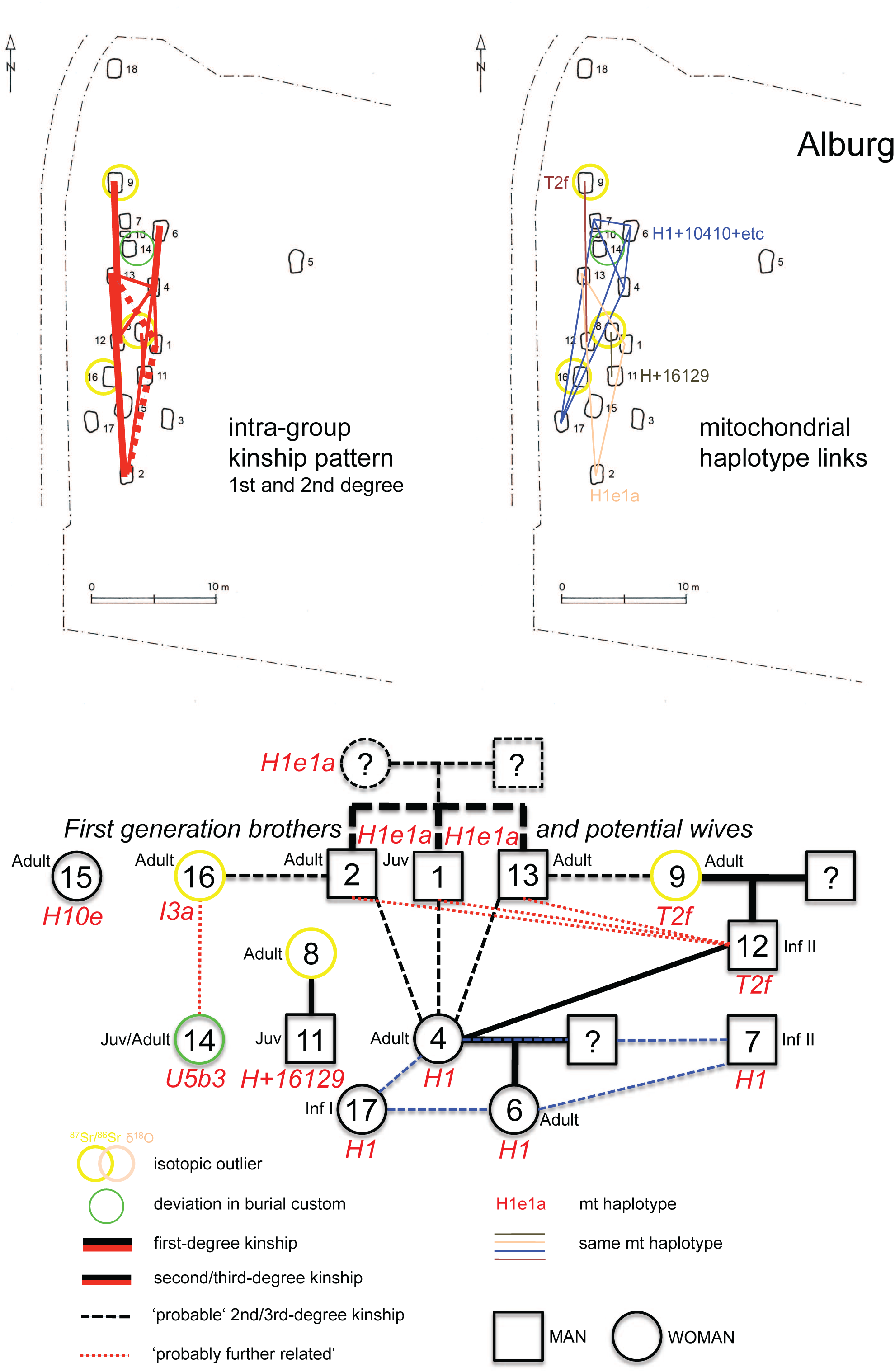
Kinship pattern indicated and genealogy reconstructed for the Alburg cemetery.

**Fig 7.**
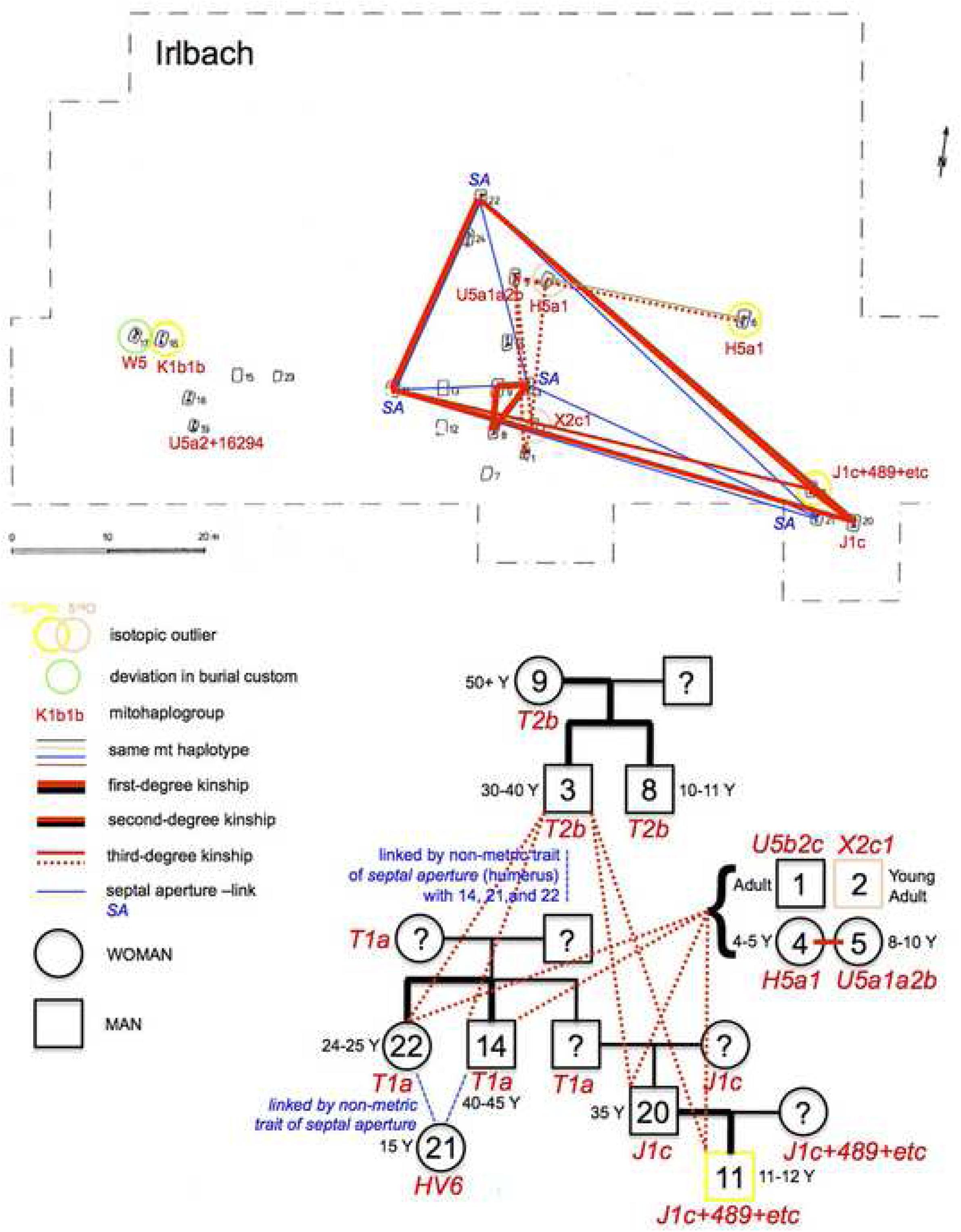
Kinship pattern indicated and genealogy reconstructed for the Irlbach cemetery.

### Isotopes

We also studied isotopic ^87^Sr/^86^Sr and δ^18^O data from tooth enamel of a total of 35 graves in the two cemeteries (19 from Irlbach: 15x ^87^Sr/^86^Sr and 18x δ^18^O; 16 from Alburg: 14x ^87^Sr/^86^Sr and 16x δ^18^O; (Fig 8). Previous studies characterized the local values of the biologically available ^87^Sr/^86^Sr for both cemeteries, located in the loess soil covered lower terraces of the right Danube bank, to be around 0.709-0.710 [19, 20, 21]. This makes the ALB 8, ALB 9 and ALB 16 and IRL 6, IRL 11 and IRL 16 ^87^Sr/^86^Sr outliers, i.e. non-locals [49]. Compared to the Bell Beaker culture burials around Augsburg, the ratio of ^87^Sr/^86^Sr values comparing non-locals to locals are nearly identical (here: 1:4.83; Augsburg: 1:4.5; 13). Of these, only the origin of ALB 16 can be geographically pinpointed due to the highly radiogenic geological background of the value 0.71740, one of the highest ever measured from Southern German samples. Its closest match is to be found just across the Danube river on the palaeozoic rocks of the *Bayerischer Wald*, the mountainous range between Bavaria and the Czech Republic. However, other more distant locations are also possible. While all Irlbach ^87^Sr/^86^Sr outliers have different values, thus likely coming from diverse geographical backgrounds, the isotopic ratios of the individuals in ALB 8 and ALB 9 are very similar, making it possible that both women, however separated by likely one or two generations, were coming from the same region and potentially community.

**Fig 8.**
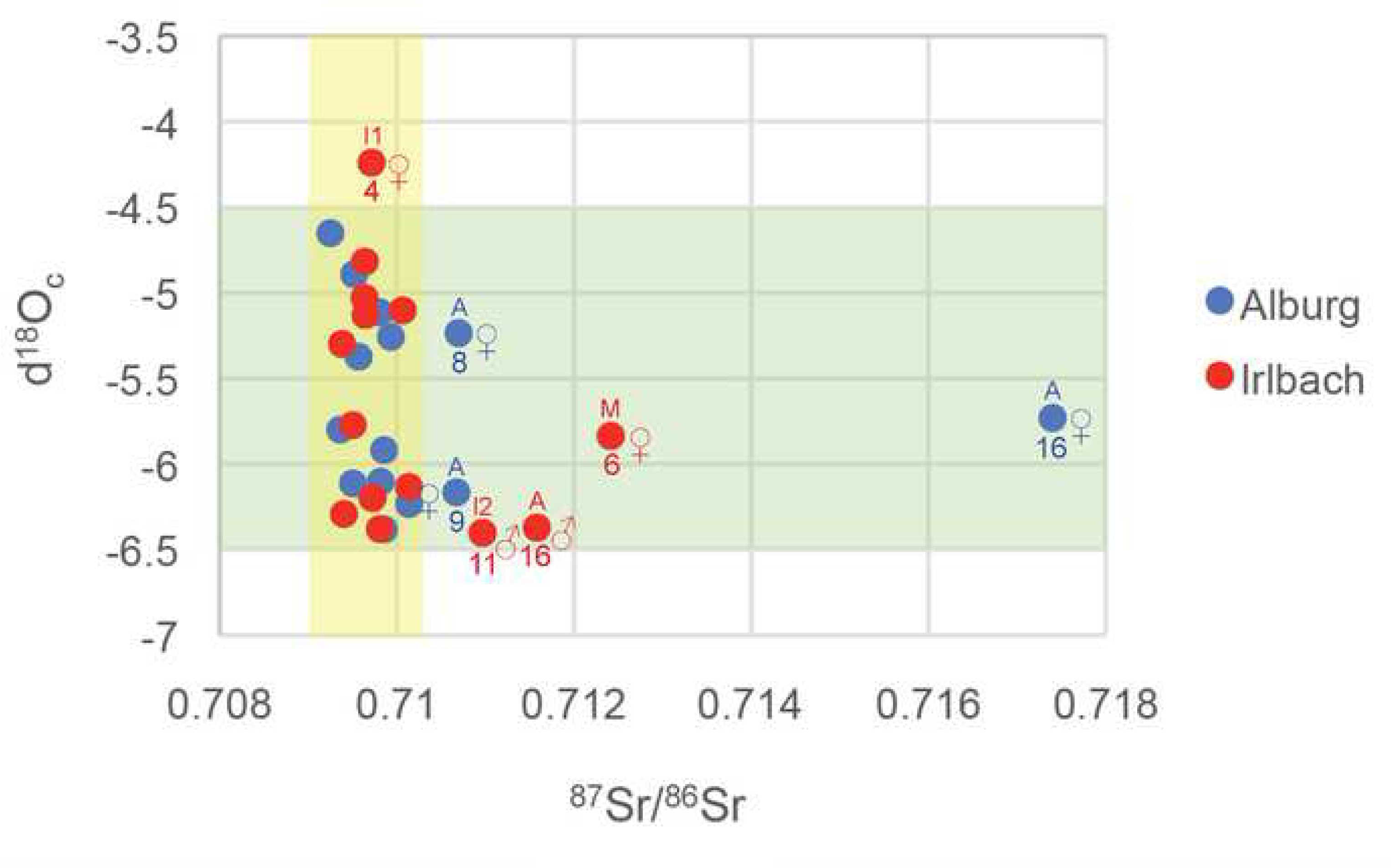
Scatter plot of ^87^Sr/^86^Sr and δ^18^Oc isotope ratios of the individuals of the Irlbach and Alburg cemeteries. Outlier graves are numbers and sex and age at death are indicated. The yellow background represents the local range of the ^87^Sr/^86^Sr ratio, the green background that of δ^18^Oc. Typical ^87^Sr/^86^Sr errors are 0.00001. Note that the δ^18^Op outlier of ALB 2 is not shown in this graph as we could not get a ^87^Sr/^86^Sr ratio for this young adult woman.

δ^18^Oc oxygen isotope data were measured in tooth enamel in structural carbonate, ‰ relative to Vienna Pee Dee Belemnite (VPDB). Local southeast Bavarian δ^18^Oc values fall into the range of −4.6 to −6.4 (= 15.0 to 17.5 when re-calculated to δ^18^Op : 13). In our dataset, two groups can be discerned, one having −4.6 to −5.4 and the other −5.75 to −6.4 values. In each group, burials from both cemeteries, both sexes, and all phases of the chronological sequence are represented. δ^18^Oc outliers are found in the IRL 2, the young adult woman of this double woman-child inhumation, and IRL 4, an infant girl aged 4-5 at death, with similar δ^18^Oc values of −4.297 and − 4.242 respectively.

Combined, eight individuals can be regarded as isotopic outliers. These represent six females (IRL 2; IRL 4; IRL 6; ALB 8; ALB 9; ALB 16) and two males (IRL 11; IRL 16). Six are adults, among which is the only man of IRL 16 from the western group. Two are children, the mentioned girl of IRL 4 and a juvenile boy, died aged 11-12, of IRL 11. None of these burials is a combined ^87^Sr/^86^Sr and δ^18^O outlier. Again, the Figs comparing non-locals to locals in Irlbach and Alburg match very well conclusions drawn from the Bell Beaker burials of the Augsburg region [13, 14] and other burial grounds in South Germany [31], consistent with two patrilocal and exogamic communities.

A more complex picture of the non-locals emerges when combining the information from archaeology, anthropology and genetics. In Alburg, two of the three isotopic outlier women belong to the first phase of the cemetery use and are buried next to the first generation brothers. However it is difficult to establish their exact relationship and how they may have been coupled due to the caveat of apparently missing family members, notably adult men (overall only three adult men versus seven women). Importantly, ALB 9, the burial with the only decorated beaker vessel in both cemeteries, appears in the PCA to be the sample with the least Steppe/Corded Ware ancestry of both cemeteries, leaning to the direction of pre-Yamnaya Neolithic populations. The burial in nearby ALB 12 is her son, passed away at age 7-14. However his father is not among the adult males we have kinship information for, although the boy is a 3rd-degree relative, much more likely a first cousin than a great grandchild, to the three brothers.

ALB 16, as said likely coming from across the Danube from the *Bayerischer Wald* region, yields the rare mitochondrial haplotype I3a and is, in contrast, among the samples with highest Steppe/Corded Ware-related ancestry from both cemeteries. In ALB 14, she has a 3rd-degree relative too, the woman placed in the grave on the body side normally reserved for men who died in juvenile/early adult age. However, ALB 14 is not at all related to the one of the three brothers buried next to ALB 16, i.e. ALB 2. They are therefore unlikely a couple unless one sees ALB 14 as another first cousin of ALB 16 and like her coming from the outside and the same family group but integrated into the Alburg community perhaps a generation later. For this conclusion could speak the fact that she shares an equally high Steppe/Corded Ware-related ancestry.

ALB 8, finally, is an adult woman who has entered the group perhaps in the second or third generation. She is 2nd-degree relative to and shares the same mitochondrial haplotype H+16129 with the juvenile boy in nearby ALB 11 and might therefore well have been his grandmother or maternal aunt. Again, we do not see the boy’s father in our records.

In Irlbach, one δ^18^O outlier is IRL 2, the young adult woman in the only double woman-child inhumation. She arrives during the middle occupation phase of the cemetery and brings in the exotic mitochrondrial haplotype X2c1. In contrast, all three ^87^Sr/^86^Sr outliers belong to the last phase of the cemetery. This could point to the arrival event of new people at this time. Here, IRL 6 is the grave of a mature (died aged 45) woman. While given an isolated place in the cemetery, she shares the same mitochondrial haplotype H5a1 and could be 3rd-degree relative with the girl IRL 4, who is one of two δ^18^O outliers.

IRL 16, an adult man, is the only outlier in the west group. He belongs to the graves with highest Steppe/Corded Ware ancestry in the PCA, and lies immediately next to the mature woman IRL 17 who, in turn, has the rare mitochondrial haplotype W5 and is the only burial of this cemetery having a deviation from the strict burial customs. They could well have been an immigrant couple but seemingly unrelated to other members of the western and central burial group. Other potential man-woman couples in this cemetery, buried next to each other however seemingly genetically unrelated, could be IRL 3 (♂) and 2 (♀) and IRL 14 (♂) and 13 (♀).

IRL 11, the juvenile son having died aged 11-12 of the neighbouring adult man IRL 20, and grand-nephew of IRL 14 and IRL 22, is also an isotopic outlier. The two form a small isolated group in the east with a third burial, IRL 21, a 15 years old girl with the mitochondrial haplotype HV6, attested only in her, and being the only girl of a marriageable age in both cemeteries. While we unfortunately have no kinship data for her, she possesses the non-metric trait of a *septal aperture* of her humerus, potentially linking her epi-genetically to the IRL 14 and IRL 22 and thus perhaps making her a relative too. All three are roughly contemporary, and may well reflect another immigrant group, with the girl perhaps to be married into the local community. However it is only the boy who has a non-local isotopic signal, different from that of his father. One possible scenario consistent with this is that his father, originally born and raised here, lived away for a while in an isotopically different environment, where his son was born and spend his youth until they returned together. Taking the multiple local relationship of his father and the possibility that he never left his group, another scenario could see the boy as a foster child, given away to a relative at an early age and returning as a juvenile shortly before passing away.

## Synthesis: expansionist kinship institutions

### Principles of social organization

By combining the various sciences, and applying them to the 42 graves of our two late Bell Beaker culture cemeteries, we propose a model characterized by six social principles:

1. The basic kinship units are nuclear families. Alburg started out with two brothers and their likely wives (a possible third brother died as teenager), which over time merged, at least genetically, into one lineage. In Irlbach we also have a family lineage as the western group is heterogeneous and unrelated to the rest. We can follow them over four to six generations, despite missing some of their members in our records, particularly adult males in the case of Alburg. Based on age distribution, these families comprised parents, some of their children of various ages, and occasionally a member of the grandparent generation. In this sense, our nuclear families are identical to those described in the Eulau (Germany) massacre, belonging culturally to the Corded Ware and being ∼400 years older [12]. They are also identical to those highlighted for the subsequent Early Bronze Age around Augsburg, mostly being 200+ years later [14].
2. These nuclear family groups are based on patriarchal, patrilinear and patrilocal residency lines (Fig 9A-B). This is exemplified by the brothers (ALB 1, ALB 12 and ALB 13) who are likely to be founders of the Alburg cemetery. It is also evident from the observed Y-chromosome homogeneity and the isotopic gender bias, as in Eulau and around Augsburg. It is further supported by the selective favouring of male child and juvenile burials (5 boys versus 3 girls; 2 juvenile boys versus 1 juvenile girl) although these Figs might be incomplete due to not having sex determinations for all 16 non-adults in the two cemeteries. It is however also observed in other South German Bell Beaker and Early Bronze Age cemeteries [45, 14]. The case of the adult woman in ALB 4 also demonstrates the important role local women of kin can have in such nuclear family groups. She is 2nd/3rd-degree relative of the three brothers, probably a grandchild/niece or great grandchild, in turn the mother of ALB 6 and in close maternal relationship with the later ALB 7 and ALB 17, thus standing in-between the generations. However due to her parent generation missing, we cannot estimate if they, or she, had spent some time away from her group.
3. The marriage system is based on female exogamy and likely monogamous. This is supported by the isotopic evidence and the equal number of male and female burials in both cemeteries, of adult men and women in Irlbach, and of missing half-siblings in our genetic records. It is also supported by the variety of mitochondrial haplotypes (Fig 3), being brought in over generations by women from various regional backgrounds. The genetic backgrounds of Irlbach and Alburg can also be quite diverse in terms of genetic lineages as shown by some exotic haplotypes and varying degrees of ancestry. Women thus came from both predominantly ‘Early/Middle Neolithic’ genetic backgrounds and predominantly ‘Steppe/Corded Ware’ genetic backgrounds. At this time they were all part of Bell Beaker culture communities. Some individuals, or groups, might even originate from down the Danube river in what is now Hungary where a higher ‘Neolithic’ genetic imprint is maintained [31, 1]. This Danube river link, more than any Únětice territory speculation, may also play a role in the increasing ‘Anatolian farmer-related ancestry’ from Bell Beaker to Early to Middle Bronze Age periods observed around Augsburg [14].
4. The inheritance system is likely based on male primogeniture. Not only are children Figs far too low in what one would expect in a prehistoric society, children are also differentially represented in burials according to age and sex/gender; the slight female deficit for infant I/II children could speak for selection and so does the juvenile gender bias for favouring boys, even if young girls could have been given away as wives. The only teenage girl in both cemeteries is likely of non-local origin, she exhibits a unique mitochondrial haplotype, and could thus be an example of a married-in girl who passed away however young at the age of 15. The possible foster boy buried next to her in IRL 11 also fits well into such a system of giving promising sons into the hands of close relatives. The practice is also observed around Augsburg in three male adults showing distinct isotopic changes in M1 and M3 teeth, resulting from seemingly having spend some time in a geological different environment before returning to their birthplace [14]. In combination with our evidence of patriarchy, patrilinearity, patrilocality, and exogamy, this likely speaks for an inheritance system along the male line and the importance of primogeniture. The latter is supported by child burials with prestigious weaponry sets, providing them with the inherited (as opposed to achieved) status of a Bell Beaker culture hunter/warrior [45].
5. Nuclear families likely formed independent households. However the question arises whether the buried family members and/or household leaders were sufficient to sustain stable households for ∼100 years, as in our cases, without having other individuals, non-family members or distant relatives, to support their households but being devoid of rights of a burial. Thus, the unequal distribution of prestige goods and hunter/warrior status in other nearby cemeteries [45: 347-352] speaks for hierarchies, ergo social inequalities, within families/households, rendering unfree and low status family members ritually invisible, in contrast to the subsequent Early Bronze Age cemeteries around Augsburg [14] and the Únětice Culture [50].
6. Families/households formed alliances through kinship and the observed exogamic practices and foster children further forged such alliances, likely linking families into clans. Alliances were thus regional rather than closely local, and they could have formed larger political and ethnic entities to be mobilized in periods of unrest, or during periods of expansion (Fig 9B). This pattern is not confined to South Germany as demonstrated by another, roughly contemporary kinship group, from the Salisbury Plain near Stonehenge in England. Here, father (I2457; 2480-2031 calBCE, 3890±30 BP, SUERC-36210; 2200-2031 calBCE, 3717±28 BP, SUERC-69975) and biological daughter (I2600; 2140-1940 calBCE, 3646±27 BP, SUERC-43374) are buried in the two different cemeteries of Amesbury Down (grave 13382; ‘adult male’) and Porton Down (grave 5108, ‘subadult female’ with a neonate), respectively, being 6.5 km apart. Two further 3rd/4th-degree male relatives of the couple are, further on, buried next to the father’s grave in Amesbury Down (I2566; grave 13385; 2210-2030 calBCE, 3734±25 BP, NZA-32490; ‘adult male’ with a long-necked beaker) and in Wilsford Down (I6777; parish of Wilsford-cum-Lake, barrow G54; a radiocarbon date is not available however seemingly richly equipped burial of a ‘17-25-year-old male’ belonging to an early Beaker phase), the latter being 3.3 kilometres away and likely a direct ancestor, while Amesbury Down, grave 13385 could be a cousin of the father/daughter couple [3]. However, no such links seem to exist between our two burial communities, as they neither share kinship, nor mitochondrial haplotypes. This picture seemingly continues into the Early Bronze Age, as shown around Augsburg [14]. There is also no exact mitochondrial match with other burials from Bavaria so far. The closest haplotypes are E09613 (haplotype H1+10410+16193) from grave 3 (feature no. 168) of the Hugo-Eckener-Straße cemetery in Augsburg, a non-local adult female, with ALB 4, ALB 6, ALB 7 and ALB 17 who share the H1+10410+16193+16286 haplotype. These haplotypes are separated by only one mutation. However their split can already have happened several generations before [13, Dateset S1, Tab. 2].

**Fig 9.**
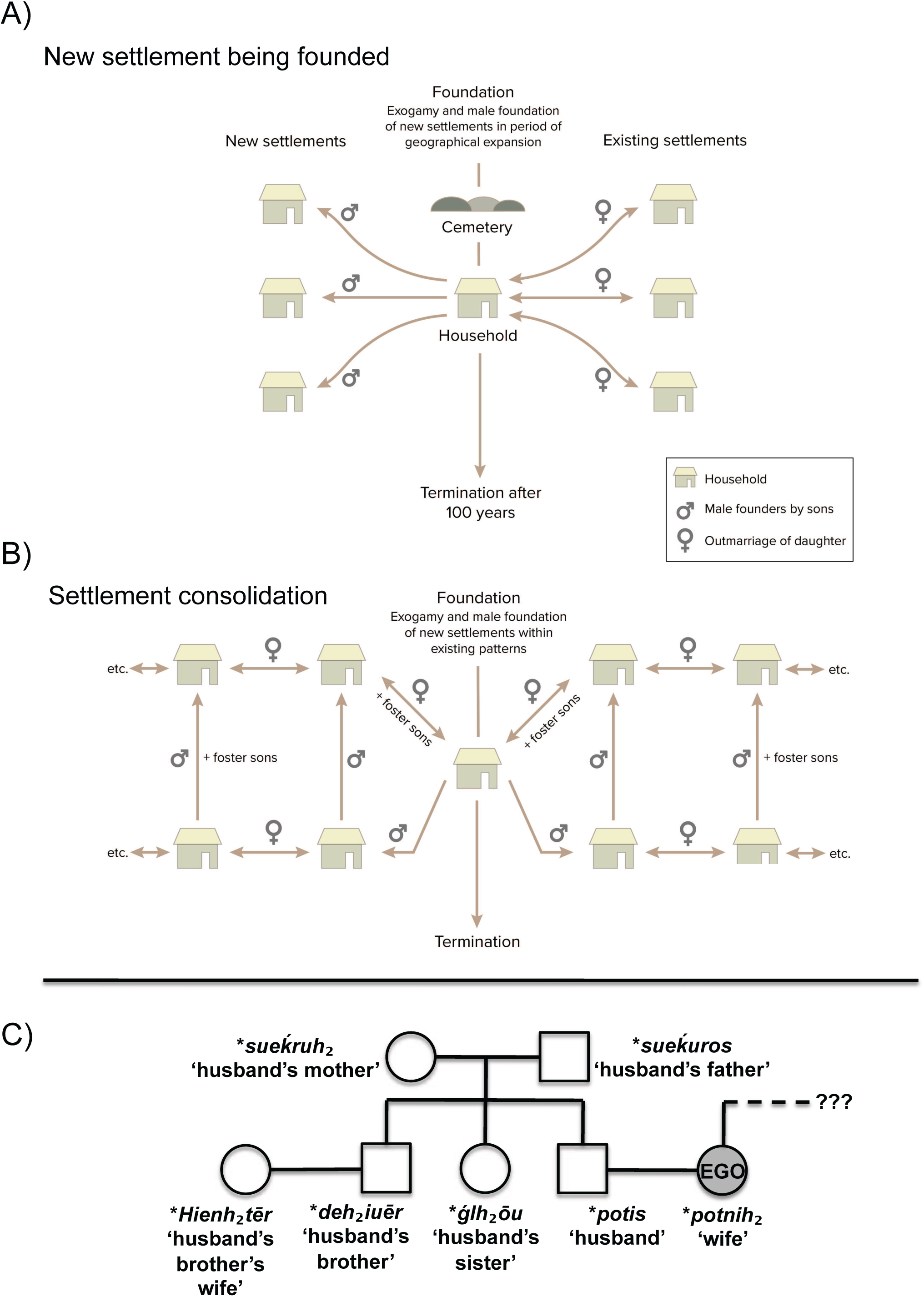
Social institutions as a model: 9A: Exogamy and male foundation of new settlements within existing patterns. 9B: Exogamy and male foundation of new settlements in period of geographical expansion. 9C: Kinship diagram of the reconstructable Proto-Indo-European terms for relatives of the marital partners. The wealth of words for relatives on the husband’s side versus the absence of those on the wife’s side is consistent with a system of patrilocal exogamy.

## Discussion

We shall now situate our results within an interpretative framework of comparative anthropology and the linguistic reconstruction of Indo-European kinship systems and institutions [51, 52].

Recent research comparing patrilocal and matrilocal marriage system in traditional societies in Indonesia demonstrated that these two institutions had different consequences for both language dominance and genetic dominance. The results were summarized as follows: “*When multiple languages are present in a region and post-marital residence rules encourage sustained directional movement between speech communities, then languages should be channelled along uniparental lines. Over time, these kinship systems shaped their gene and language phylogenies*” [53]. Consequently, women marrying into patrilocal communities were forced to adopt their husband’s language. Such a situation could well resemble third millennium BC Europe. If Indo-European speakers were the ones to introduce a system of patrilocal exogamy, women from one or multiple originally non-Indo-European-speaking communities would have moved into Indo-European speaking communities, and adopt their language [6]. Over time this would lead to an increased genetic and cultural dominance and the consolidation of one or more Indo-European dialects.

Our data allows us to identify the institution of exogamy linked to patrilocal/virilocal residence for women. These results are supported by the identification of a father and daughter buried in British Bell Beakers context in Amesbury Down and Porton Down, being 6.5 km apart. This kinship model was widespread among historical Indo-European-speaking societies and has previously been hypothesized for the Proto-Indo-European language (PIE) community by methods of linguistic reconstruction [52: 212, 54, 55, 56, 57, 58]. Linguistic indicators of exogamy consist mainly of reconstructed Proto-Indo-European vocabulary such as a word for ‘brideprice’ (*h1uedmōn), and the synonymy of the verbs ‘to wed’ and ‘to lead’ (*uod^h^eieti), suggesting that the bride was led away from her ancestral to her new husband’s household [55: 199). Patrilocality and the consequential remoteness of the wife’s relatives is further implied by the fact that the Proto-Indo-European reconstructed kinship terms show a strong bias towards names for the relatives of the husband as opposed to a marked absence for those of the wife (Fig 9C). Our study thus provides the first identification of potential continuity between the linguistically reconstructed kinship structures of Proto-Indo-European speakers in the late third millennium BC and that of their linguistic descendants as they are found in the earliest historical sources. In our data, we are further able to identify one dominant genetic male lineage, while there are multiple female lines, suggesting a strong patriarchal and patrilineal dominance through time. This resembles the patrilineal household that has been reconstructed for Proto-Indo-European, consisting of the master of the house (*dems potis), his wife (*potnih2), sons (*suHnus), unmarried daughters (*d^h^ugh2tēr), daughters-in-law (*snusos) and grandchildren (*nepotes) [52: table 12.1, 54, 55, 58, 59]. This model bears resemblance to the so-called Omaha kinship system. The Omaha kinship system can be characterized in the following way: “*Agnatic ties, especially between male siblings are emphasised, and the household made up of agnatically related males, their wives and offspring, is usually highly solidary and the most important political and economic unit. Strong controls are exercised over the actions of group members, usually under the autocratic rule of the household head, and particularly over wives and their offspring. Residence at marriage is strictly virilocal, bride wealth payments are usually high and there may be severe sanctions against divorce and adultery*” [56]. Accordingly, our two cemeteries represent household leaders and their close kin.

In the Omaha kinship system those male lineages/households that were successful in marrying out their daughters to alliance partners ‘would not only receive more bride wealth than others, but also have the potential for receiving foster sons who would move to their mother’s brother and become young warriors’ [60: 238]. Fosterage of young boys in their mother’s family was common in early Indo-European-speaking societies, such as Germanic and Celtic groups, typically at a maternal uncle [57, 58, 61, 62: 123]. We identified one possible case in Irlbach, the juvenile boy IRL 11 displaying a non-local ^87^Sr/^86^Sr signature, whose genetic father lay next to him in the cemetery and had a different and local signature. Thus, this boy could had been raised in a different locality, perhaps with his maternal uncle, who in *Omaha* terminology is equated with the grandfather (PIE *h2euh2os), and then returned just before entering adolescent age. Similar evidence is perhaps documented for Eulau, grave 98 where the supposed mother of the two children in this triple burial is definitely not the biological mother [10: 18228], and for seemingly three burials around Augsburg [14, 63: 255]. It corresponds to the observation that the word for ‘brother’ in early Indo-European was used in a wider sense, to indicate a group of young males related by kinship or common social affiliation, e.g. members of the same ‘brotherhood’ [52: 214], an institution also documented archaeologically [64].

An important aspect of the *Omaha* kinship system is its flexibility and potential for expansion. Although highly opportunistic, it had one rule that one was not allowed to marry twice into the same family. If such a rule was in place, this would in turn produce more variable alliances with other families, and thus expand potential political support, whether for exchange/trade or when mobilizing for hunt/warfare. In accordance with this, we can identify multiple female mitochondrial haplotypes at both cemeteries, identifying female genetic inheritance. Thus, while there is only one dominant Y-chromosome haplogroup (R1b-M269-P312), there are multiple female lineages, and not a single mitochondrial haplotype is identical in both cemeteries despite them being only 17 kilometres apart and mostly contemporaneous. This evidence supports the notion that marriage was indeed an instrument in creating widespread alliances, which was useful in a settlement structure of singular homesteads spread out in the landscape. It also supports the proposition by Knipper et al. [13] that genetic diversity increases over time in such a system.

While the linguistic reconstruction of the original Proto-Indo-European language system roughly corresponds with an Omaha system, the traditions of the Bell Beaker communities were no doubt characterized by innovations. For instance, the typically Omaha feature called generational skewing, i.e. the use of identical kinship terms for male cross-relatives on the mother’s side, cannot be reconstructed for the Indo-European proto-language [contra 65]. The evidence for this feature emerged independently in several Indo-European languages spoken in Europe, and is absent in Asia [66]. It therefore likely evolved secondarily in post-Yamnaya contexts. When mobile steppe pastoralists adopted a more sedentary lifestyle, allowing for intensified contact with other nuclear families and especially relatives of the mother, new kinship terms were added to the languages along with new kinship roles. It is this innovated patrilineal and patrilocal kinship model that could have facilitated the spread of Indo-European dialects according to the model proposed by Lansing et al. [53].

Our cemeteries are slightly later than those Bell Beaker grave groups around Augsburg, with some good overlap existing, and the diversity of mitochondrial haplotypes in Irlbach is similar to those in Augsburg, while is lower in Alburg. Rather it seems that higher or lower number of mitochondrial haplotypes reflect either group hetero/homogeneity or successful or less successful marriage strategies. Regarding the latter, it seems that competition was at work between the settlements in Irlbach and Alburg as they do not share any mitochondrial female lines. This raises the question how far-reaching such marriage alliances were during the Bell Beaker culture and its time. We have in Alburg one case of a woman coming from across the Danube river, and the other ^87^Sr/^86^Sr outliers perhaps from the same place. Around Augsburg, Knipper et al. 2017 assumes a catchment along the Lech river, with the nearby Ries region, c. 60km away, being the nearest occurrence of more radiogenic ^87^Sr/^86^Sr values. We now know that during the Nordic Bronze Age, a 1000 years later, young women could move 4-500 kilometres or more, as in the case of the ‘Skrydstrup woman’ [67], if the interpretation of the bio-available strontium for that site is correct [68].

Gender balance is nearly one to one between males and females. Having also failed to detect half-siblings, our evidence thus suggests monogamy as a dominant principle. When it comes to age differentiation there are some imbalances: There are clearly more juvenile (teenage) men (5:1), which suggests higher mortality or perhaps selection of certain males for burial. Here we should consider the effects of primogeniture, which implied strong continuity in the transmission of property as well as genes in the male line. However, it also produces males who would have to look elsewhere for their future. These are therefore a strong mobilizing group for colonizing new settlements and thus expand the group, but until they are initiated as grown up they are often organized in a special institution of youth war bands to train them for their future [6, 41]. This may also have been a period of risks and early death, whereas young girls were probably married out already when they entered puberty, which might go some way to explain the discrepancy in the cemetery. This reminds again of the later cases around Augsburg [14] and of the ‘Skydstrup woman’, who had moved from Central Europe to Denmark at the age of 14, most probably as part of a marriage alliance [67].

Our two cemeteries not only started but also terminated at approximately the same time, which corresponds to periods of wider termination of cemeteries in the region and the foundation of cemeteries in new locations [15, 45, 31]. There seems thus to have existed a certain dynamic in the settlement system, with cycles of changing locations after ∼100 years. At Alburg, the first to be buried were two brothers, with perhaps a third brother to have been buried next to them. However only the two lived long enough to became fathers and grandfathers of later offspring to be also buried in the cemetery. Three brothers as founding fathers play a special role in much later Indo-European folklore and mythology [69], and may reflect the role of sons without primogeniture inheritance in founding new families/households. It corresponds to the role of triplism in mythology and symbolism [70: 208]. While we have documented that the cemeteries contained four to six family generations, members of new and of related kin appear to have joined the existing group at Irlbach towards the end of the cycle, perhaps indicating the start of the relocation of the settlement and a new period of expansion.

These observations underline the inherent expansionist dynamics of the social system we have described with continuity from Corded Ware to Bell Beaker culture. This type of social organization stands, however, in some opposition to genetic and ^87^Sr/^86^Sr results from the Globular Amphora culture mass burial of Koszyce (Poland), being ∼600 years earlier [71]. Similar to our example, mitochondrial variation (six haplotypes) is larger than Y-chromosome variation, as only one Y chromosome haplotype has been identified. This suggests exogamic marriage relations, virilocal residence and patrilineal descent, like in Alburg and Irlbach, and probably widely practised in Neolithic [72] and Early Bronze Age [13] Europe. However, there are also differences. At Koszyce it seems that four nuclear families, or parts of them, form a single large extended family and therefore the kinship principles might be of a different nature. The mitochondrial haplotypes are far less varied than in the Bell Beaker culture (six different ones out of 15 individuals) and several brothers were sharing the same father but different mothers, who in turn might have nevertheless been related to each other [71]. This speaks either of a non-monogamous system, or of serial monogamy. Also, the low mitochondrial variation can suggest a fundamentally different marriage system, or is just an indication of an overall more genetically homogenous society. Thus, there are indications that Corded Ware and Bell Beaker social organization was of a different nature than that of preceding Neolithic societies, but there is still much to be learned from future research.

## Conclusions

The extraction and combination of many types of evidence –archaeological, anthropological, strontium/oxygen isotopes and ancient DNA– has allowed an unprecedented high-resolution interpretative narrative of the lives of two families who lived little more than 4000 years ago. The results correspond to what is known about the earliest attested Indo-European societies and the linguistic reconstruction of the Indo-European proto-language. The evidence sustained the reconstruction of a kinship structure that was based on a dominant male line that married in women from other groups and married out their own daughters in this way building up a network of alliances, which could become part of a competitive mobilization in periods of unrest, and perhaps also to secure access to resources like metal. Such a system corresponds to the well-documented *Omaha* kinship structure of exogamy linked to virilocal residence and patrilineal descent, and primogeniture. It favored an expansive settlement policy, and it also provides for the first time a realistic model for the spread of a prehistoric language family along male lineages, as well as their genetic dominance through time. As a result most modern Europeans of the northern and western half of the Continent are genetically related to Corded Ware and Bell Beaker people of the third millennium BC, and probably speak evolved forms of their languages as well. It should, however, be noted that although archaeological and linguistic evidence provide a rather homologous or shared picture, such a social organization is not exclusively Indo-European but may also be found in later pastoral / agro-pastoral societies, based on similarities in economic organization [73], also reflected in the *Omaha* kinship system. Thus, we propose that our archaeological case study provides a historically particular match between archaeogenetic and linguistic-anthropological reconstructions of the kinship systems that may attain stronger generalizing power by future studies [4]. However, since our results concur with those from the Lech valley near Augsburg in Bavaria, it seems likely that we are dealing with an institutionalized practice covering a wider segment of Bell Beaker society, however rooted in Corded Ware society. Our model of an expansionist social organization may thus serve as a test case for further comparative studies.

## Acknowledgements

Francois Bertemes, Richard Harrison, Kate Robson-Brown and Volker Heyd wish to thank the Fritz-Thyssen-Stiftung in Germany for its generous support of the then project RR8322 -- Kinship and Residence Patterns in the Late Copper Age of Southern Germany -- from which this research originally stems. Kristian Kristiansen and Karl-Göran Sjögren were funded by the Swedish Riksbanken grant M16-0455:1 -- Towards a New Prehistory. The research here forms also part of Volker Heyd’s ERC Advanced project 788616: The Yamnaya Impact on Prehistoric Europe (YMPACT). We also want to thank Eske Willerslev, Geogenetics, Copenhagen, for his support of this research. David Reich wishes to mention that he is also an investigator of the Howard Hughes Medical Institute. We are grateful to Tian Chen Zeng, Dept of Human Evolutionary Biology at Harvard University, for his work on better visualizing Fig 5.

Supporting information table. Sample details.

## References

Allentoft ME et al., Population genomics of Bronze Age Eurasia. Nature 2015; 522: 167–172. Doi: 10.1038/nature14507.

Haak W et al., Massive migration from the steppe was a source for Indo-European languages in Europe. Nature 2015; 522: 207–211. Doi: 10.1038/nature14317.

Olalde I et al., The Beaker Phenomenon and the Genomic Transformation of Northwest Europe. Nature 2018; 555: 190–196. Doi: 10.1038/nature25738.

Furholt M, Re-integrating Archaeology: A Contribution to aDNA Studies and the Migration Discourse on the 3rd Millennium BC in Europe. Proc Prehistoric Soc 2019; 85. Doi: 10.1017/ppr.2019.4.

Iversen R, Kroonen G, Talking Neolithic: Linguistic and Archaeological Perspectives on How Indo-European Was Implemented in Southern Scandinavia. American J Arch 2017; 121/4: 11–25. Doi: 10.3764/aja.121.4.0511.

Kristiansen K et al., Re-theorising mobility and the formation of culture and language among the Corded Ware Culture in Europe. Antiquity 2017; 91: 334–347.

Sjögren K-G, Price TD, Kristiansen K, Diet and mobility in the Corded Ware of central Europe. PLoS One 2016; 11:e0155083.

Goldberg A, Günther T, Rosenberg NA, Jakobsson M, Ancient X chromosomes reveal contrasting sex bias in Neolithic and Bronze Age Eurasian migrations. Proc Natl Acad Sci USA 2017; 114/10: 2657–2662.

Zeng TC, Aw AJ, Feldman MW, Cultural hitchhiking and competition between patrilineal kin groups explain the post-Neolithic Y-chromosome bottleneck. Nature Comm. 2018; 9: 2077. Doi: 10.1038/s41467-018-04375-6.

Lazaridis I, Reich D, Failure to replicate a genetic signal for sex bias in the steppe migration into central Europe. Proc Natl Acad Sci USA 2017; 114/20: E3873– E3874. Doi: 10.1073/pnas.1704308114

Goldberg A, Günther T, Rosenberg NA, Jakobsson M, Reply to Lazaridis and Reich: Robust model-based inference of male-biased admixture during Bronze Age migration from the Pontic-Caspian Steppe. Proc Natl Acad Sci USA 2017; 114/20: E3875– E3877. Doi: 10.1073/pnas.1704442114

Haak W, et al., Ancient DNA, Strontium isotopes, and osteological analyses shed light on social and kinship organization of the Later Stone Age. Proc Natl Acad Sci USA 2008; 105: 18226–18231.

Knipper C et al., Female exogamy and gene pool diversification at the transition from the Final Neolithic to the Early Bronze Age in central Europe. Proc Natl Acad Sci USA 2017; 114: 10083–10088. Doi: 10.1073/pnas.1706355114

Mittnik A et al., Kinship-based social inequality in Bronze Age Europe. Science 2019; Doi: 10.1126/science.aax6219.

Heyd V, Die Spätkupferzeit in Süddeutschland. Habelt; 2000.

Brothwell DR, Digging up Bones, 3rd edition. Cornell University Press; 1981.

White TD, Folkens PA, The Human Bone Manual. Elsevier Academic Press; 2005.

Bass WM, Human Osteology: a field and laboratory guide. Special Publication No. 2 of the Missouri Archaeological Society. Springfield; 1987.

Chamberlain A, Interpreting the Past: Human Remains. British Museum Press; 1994.

Scheuer L, Black S, Developmental Juvenile Osteology. Academic Press; 2000.

Meindl RS, Lovejoy CO, Ectocranial Suture Closure: A Revised Method for the Determination of Skeletal Age at Death Based on the Lateral-Anterior Sutures. American J Physical Anthrop 1985; 68: 57–66.

Berry AC, Berry RJ, Epigenetic variation in the human cranium. J Anatomy 1967; 101: 361–379.

Buikstra JC, Ubelaker DH, Standards for Data Collection from Human Skeletal Remains. Arkansas Archaeological Survey, Research Series 44. Fayetteville; 1994.

Mann RW, Hunt DR, Lozanoff S, Photographic Regional Atlas of Non-Metric Traits and Anatomical Variants in the Human Skeleton. Charles C Thomas Publisher Ltd; 2016.

Hauser D, De Stefano GF, Epigenetic Variants of the Human Skull. Schweizerbart Science Publishers; 1989.

Trotter M, Estimation of stature from intact long limb bones. In: Stewart TD, ed. Personal Identification in Mass Disasters. National Museum of Natural History Washington; 1970. pp. 71–84.

McKinley JI, Compiling a skeletal inventory: disarticulated and co-mingled remains. In: Brickley M, McKinley JI, eds. Guidelines to the Standards for Recording Human Remains. Institute of Field Archaeologists paper 7; 2004. pp. 14–17.

Winterholler B, Untersuchungen zur Mobilität archäologischer Gruppen anhand von Strontiumisotopenverhältnissen. Unpublished diploma thesis. TU Freiburg; 2004.

Price TD, Knipper C, Grupe G, Smrcka V, Strontium isotopes and prehistoric human migration. Eur J Arch 2004; 7: 9–40.

Heyd V, Winterholler B, Böhm K, Pernicka E, Mobilität, Strontiumisotopie und Subsistenz in der süddeutschen Glockenbecherkultur. Ber Bayer Bodendenkmalpfl 2004; 43/44: 109–135.

Bertemes F, Heyd V, 2200 BC – Innovation or Evolution? In: Meller H et al. eds. 2200 BC – A Climatic Breakdown as a Cause for the Collapse of the Old World? Landesamt für Denkmalpflege und Archäologie Sachsen-Anhalt; 2015. pp. 561–578.

Steiger RH, Jaeger E Subcommission of geochronology – convention on the use of decay constants in geo- and cosmo-chronology. Earth and Planetary Sci Letters 1977; 36: 359–362.

Rohland N, Harney E, Mallick S, Nordenfelt S, Reich D, Partial Uracil – DNA – Glycosylase Treatment for Screening of Ancient DNA. Philosophical Transactions R Soc London B 370, 2015. Doi: 10.1098/rstb.2013.0624.

Fu Q et al., An Early Modern Human from Romania with a Recent Neanderthal Ancestor. Nature 2015; 524: 216–219. Doi: 10.1038/nature14558.

Weissensteiner H et al., HaploGrep 2: Mitochondrial Haplogroup Classification in the Era of High-Throughput Sequencing.” Nucleic Acids Res 2016; 44 (W1): W58–63. Doi: 10.1093/nar/gkw233.

Skoglund P, Storå J, Götherström A, Jakobsson M, Accurate Sex Identification of Ancient Human Remains Using DNA Shotgun Sequencing. J Arch Sci 2013; 40/12: 4477–4482. Doi: 10.1016/j.jas.2013.07.004.

Olalde I et al., The Genomic History of the Iberian Peninsula over the Past 8000 Years. Science 2019; 363: 1230–1234.

Patterson N, Price AL, Reich D, Population Structure and Eigenanalysis. PLoS Genetics 2006; 2/12: e190 (2006). Doi: 10.1371/journal.pgen.0020190.

Mathieson I et. al., Genome-Wide Patterns of Selection in 230 Ancient Eurasians. Nature 2015; 528: 499–503. Doi: 10.1038/nature16152.

Lipson M et al., Parallel Ancient Genomic Transects Reveal Complex Population History of Early European Farmers. Nature 2017; 551: 368–372. Doi: 10.1101/114488.

Cassidy L et al., Neolithic and Bronze Age Migration to Ireland and Establishment of the Insular Atlantic Genome. Proc Natl Acad Sci USA 2016; 113/2: 1–6. Doi: 10.1073/pnas.1518445113.

Olalde I et al., A Common Genetic Origin for Early Farmers from Mediterranean Cardial and Central European LBK Cultures. Molecular Biology and Evolution 2015; 32: 3132–3142. Doi: 10.1093/molbev/msv181.

Kilinc G et al., The Demographic Development of the First Farmers in Anatolia. Current Biology 2016; 26: 1–8. Doi: 10.1016/j.cub.2016.07.057.

Omrak A, Genomic Evidence Establishes Anatolia as the Source of the European Neolithic Gene Pool. Current Biology 2016; 26: 270–275. Doi: 10.1016/j.cub.2015.12.019.

Heyd V, Families, Prestige Goods, Warriors and Complex Societies: Beaker Groups of the 3rd Millennium cal BC along the Upper and Middle Danube. Proc Prehistoric Soc 2007; 73: 321–370.

Pozzi L, Ramiro Farinas D, Infant and child mortality in the past. Ann démographie historique 2015; 129/1: 55–75.

Rebay-Salisbury K. et. al., Motherhood at Early Bronze Age Unterhautzenthal, Lower Austria. Arch Austriaca 2018; 102: 71–134.

Sahaipal DT, Pichora D, Septal aperture: an anatomic variant predisposing to bilateral low-energy fractures of the distal humerus. Canadian J Surgery 2006; 49: 363–364.

Price TD, Grupe G, Schröter P, Reconstruction of migration patterns in the Bell Beaker period by stable strontium isotope analysis. Appl Geochem 1994; 9: 413–417.

Knipper C, A Distinct Section of the Early Bronze Age Society? Stable Isotope Investigations of Burials in Settlement Pits and Multiple Inhumations of the Unetice Culture in Central Germany. American J Physical Anthrop 2016; 159/3: 496–516. Doi: 10.1002/ajpa.22892.

Johnson KM, Paul KS, Bioarchaeology and Kinship: Integrating Theory, Social Relatedness, and Biology in Ancient Family Research. J Arch Res 2016; 24: 75–123.

Mallory JP, Adams DQ, The Oxford Introduction to Proto-Indo-European and the Proto-Indo-European World. Oxford University Press; 2006.

Lansing JS et al., Kinship structures create persistent channels for language transmissions. Proc Natl Acad Sci USA 2017; 114/49: 12910–12915. Doi: 10.1073/pnas.1706416114

Delbrück B, Die indogermanischen Verwandschaftsnamen: ein Beitrag zur vergleichenden Alterthumskunde. Leipzig; 1889.

Szemerényi O, Studies in the kinship terminology of the Indo-European languages, with special reference to Indian, Iranian, Greek and Latin. Acta Iranica 17. Brill; 1977.

Rowlands M, Kinship, alliance and exchange in the European Bronze Age. In: Barrett J, Bradley R, eds. Settlement and Society in the British Later Bronze Age. BAR British Series 83. Archaeopress; 1980. pp. 15–55.

Olsen BA, Kin, clan and community in Proto-Indo-European. In: Whitehead Nielsen B, Olsen BA eds. Kin, clan and community in prehistoric Europe Museum. Tusculanum Press; 2019.

Olsen BA, Aspects of family structure among the Indo-Europeans. In: Olsen BA, Olander T, Kristiansen K, eds. Indo-European languages and society in prehistory. Oxbow Books; 2019. pp. 145–164.

Pronk TC, On mobility, kinship and marriage in Indo-European society (forthcoming).

Kristiansen K, Larsson TB, The rise of Bronze Age society. Travels, Transmissions and Transformations. Cambridge University Press; 2005.

Bremmer J, Avunculate and fosterage. J Indo-European Stud 1976; 4: 65–67.

Mallory JP, In search of the Indo-Europeans: language, archaeology, and myth. Thames and Hudson; 1989.

Massy K, et al., Patterns of Transformation from the Final Neolithic to the Early Bronze Age: A Case Study from the Lech Valley South of Augsburg. In: Stockhammer PW, Maran J, eds. Appropriating innovations. Entangled knowledge in Eurasia, 5000–1500 BCE. Oxbow; 2017. pp. 241–261.

Anthony DW, Brown D, The dogs of war: A Bronze Age initiation ritual in the Russian steppes. J Anthrop Arch 2017; 48: 134–148.

Friedrich P, Proto-Indo-European Kinship. Ethnology 1966; 5: 1–36.

Hettrich H, Indo-European Kinship Terminology in Linguistics and Anthropology. Anthrop Linguistics 1985; 27: 453–480.

Frei KM et al., A matter of months: High precision migration chronology of a Bronze Age female, PLoS One 2017; 12(6): e0178834.

Thomsen E, Andreasen R, Agricultural lime disturbs natural strontium isotope variations: Implications for provenance and migration studies. Science Advances 2019; 5: eaav8083.

Graça da Silva S, Tehrani JJ, Comparative phylogenetic analyses uncover the ancient roots of Indo-European folktales. R Soc Open Sci 2016; 3: 150645. Doi: 10.1098/rsos.150645.

Green M, The Gods of the Celts. Sutton; 1986.

Schroeder H et al., Blood ties: Unravelling ancestry and kinship in a Late Neolithic mass grave. Proc Natl Acad Sci USA 2019; 116/22: 10705–10710. Doi: 10.1073/pnas.1820210116.

Brown K A, Women on the move: the DNA evidence for female mobility and exogamy in prehistory. In: Leary J ed. Past Mobilities; Archaeological Approaches to Movement and Mobility. Ashgate Publishing; 2014. pp. 155–173.

Lincoln B, Priests, Warriors and Cattle. A Study in the Ecology of Religion. University of California Press; 1981.

